# Information Decomposition in the Frequency Domain: a New Framework to Study Cardiovascular and Cardiorespiratory Oscillations

**DOI:** 10.1101/2020.10.14.338939

**Authors:** Luca Faes, Riccardo Pernice, Gorana Mijatovic, Yuri Antonacci, Jana Cernanova Krohova, Michal Javorka, Alberto Porta

## Abstract

While cross-spectral and information-theoretic approaches are widely used for the multivariate analysis of physiological time series, their combined utilization is far less developed in the literature. This study introduces a framework for the spectral decomposition of multivariate information measures, which provides frequency-specific quantifications of the information shared between a target and two source time series and of its expansion into amounts related to how the sources contribute to the target dynamics with unique, redundant and synergistic information. The framework is illustrated in simulations of linearly interacting stochastic processes, showing how it allows to retrieve amounts of information shared by the processes within specific frequency bands which are otherwise not detectable by time-domain information measures, as well as coupling features which are not detectable by spectral measures. Then, it is applied to the time series of heart period, systolic and diastolic arterial pressure and respiration variability measured in healthy subjects monitored in the resting supine position and during head-up tilt. We show that the spectral measures of unique, redundant and synergistic information shared by these variability series, integrated within specific frequency bands of physiological interest, reflect the mechanisms of short term regulation of cardiovascular and cardiorespiratory oscillations and their alterations induced by the postural stress.

## 1. Introduction

The spontaneous oscillations exhibited by the physiological variables that describe the dynamic activity of the heart, cardiovascular and respiratory systems are the focus of an intense interdisciplinary research since more than three decades. In particular, the beat-to-beat variability of the heart period (HP), systolic and diastolic arterial pressure (SAP, DAP) and respiratory volume or flow (RESP) are of great physiological relevance as they reflect the short-term cardiovascular and cardiorespiratory regulation [1]. Such regulation results from the complex interplay of several physiological mechanisms of mechanical origin and/or mediated by the sympathetic and parasympathetic branches of the autonomic nervous system (ANS) [2]. The main investigated mechanisms are the arterial baroreflex whereby SAP and DAP changes induce changes in HP aimed to maintain pressure homeostasis [3,4], the Starling law and arterial Windkessel effect whereby variations of HP affect DAP and SAP through modifications of the end diastolic volume, of the systolic contraction strength, and of the diastolic blood pressure decay duration [5,6], and the influence of RESP on SAP and HP related to mechanical effects on intrathoracic pressure and stroke volume and to alterations of the central vagal outflow exerted by respiratory neuron firing [7,8]. These diverse mechanisms interact and even compete with each other to accomplish the homeostatic control of the physiological variables. Moreover, the degree of involvement of these mechanisms into the cardiovascular regulation changes in response to alterations of the physiological state such as those induced by different types of stress [6,9–13].

The mechanisms responsible for the short-term cardiovascular and cardiorespiratory control can be probed non-invasively applying tools for multivariate time series analysis to the spontaneous variability of HP, SAP, DAP and RESP [14,15]. In this context, tools taken from the field of information theory are increasingly employed for the multivariate analysis of physiological time series [16]. The framework of information dynamics provides a versatile and unifying set of measures which allow to quantify, from multivariate time series describing the dynamic activity of multiple systems, the amounts of information produced and stored in each system, transferred from a ‘source’ system to a ‘target’ system, and modified as a consequence of the interaction between source systems sending information to a target [16–18]. These approaches are probabilistic and thus inherently model-free, but often their computation resides on linear parametric models whereby concepts of predictability and information storage, and Granger causality and information transfer, have been related [16,19,20]. With regard to physiological variability, the concept of information transfer can be used to assess the functional mechanisms underlying the coupling between two variables, while information modification allows to investigate the nature of the interactions among multiple variables. In particular, two sources are redundant if each carries individually information about the same aspects of the target, while synergy arises from independent mechanisms of interaction between each source and the target. These concepts, implemented in the measures of interaction information [18,21] and partial information decomposition (PID) [22,23], have been successfully used to investigate the mechanisms of cardiovascular and cardiorespiratory interaction in a variety of pathophysiological states [6,11,16,24–26].

A drawback of information dynamic measures is that they are not frequency-specific, meaning that they evaluate the “overall” interaction among multiple time series without focusing on specific rhythms. On the other hand, heart rate, blood pressure and respiratory variability series display a rich oscillatory content, which is typically manifested within the so-called low frequency (LF, 0.04-0.15 Hz) and high frequency (HF, 0.15-0.4 Hz) bands [1,2]. The frequency domain analysis of cardiovascular and cardiorespiratory oscillations is commonly performed using spectral measures of coupling and causality, such as the coherence and partial coherence or the directed coherence and partial directed coherence [14,27]. Nevertheless, tools for the spectral decomposition of multivariate information measures, able to provide a frequency-specific quantification not only of bivariate couplings but also of higher order interactions like the interaction information or the redundancy and synergy measures elicited by PID, are still lacking in the field of time series analysis. To fill this gap, the present work proposes a new framework for the spectral analysis of the information shared between a target process and two source processes, and for its decomposition into amounts related to how the sources contribute unique, redundant and synergistic information to the target. All measures are derived intuitively from the cross-spectral matrix of the three interacting processes, and are first tested in a theoretical example showing their behaviour in controlled conditions of multivariate coupling. Then, the framework is applied to the HP, SAP, DAP and RESP series measures from healthy subjects monitored in the supine and upright body positions in order to assess, separately for LF and HF oscillations, the mechanisms underlying cardiovascular regulation (series HP, SAP, DAP) and cardiorespiratory regulation (series HP, SAP, RESP) in a resting state and in response to the alterations in the sympatho-vagal balance induced by the postural stress.

## 2. Framework for the spectral decomposition of information dynamics

In this section we first review the measures typically used to assess dynamic interactions between two stochastic processes (i.e., interactions based on time-lagged dependencies) in the frequency domain, also evidencing the link between these measures and dynamic information quantities. Then we extend the formulation to the multivariate case, defining measures that quantify how the dynamic coupling between a “target” process and two “source” processes changes comparing collective interactions (i.e., interactions measured when the sources are grouped together) and individual interactions (i.e., interactions measured when the sources are kept separate). These higher-order interactions are evaluated in both the frequency and information-theoretic domains, providing a link between the two domains. The frequency domain information measures presented in this work are collected in the fdPID Matlab toolbox, which is publicly available at www.lucafaes.net/fdPID.html.

### 2.1 Spectral and Information-Theoretic Coupling Measures in Bivariate Processes

Let us consider two discrete-time zero mean stationary stochastic processes 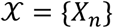 and 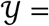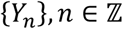 typically obtained sampling continuous-time processes *X*_*t*_ and *Y*_*t*_, *t* ∈ ℝ, at times 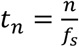 where *f*_*s*_ is the sampling frequency. In a linear analysis framework, the dynamic analysis of the interactions between the processes is typically performed in the time domain relating the variable sampling the present state of one of the two processes, say *X*_*n*_, to the variables sampling the past states of the other process, {*Y*_*n−k*_: *k* ≥ 0}, through the cross-correlation function 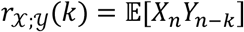.

Moving to the frequency domain, a normalized measure of coupling between 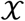 and 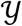 is obtained considering the power spectral density (PSD) of each process defined as the Fourier Transform (FT) of its autocorrelation function (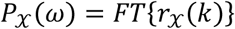, 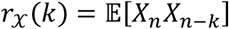 and the same for 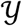), and their cross PSD defined as the FT of the cross-correlation 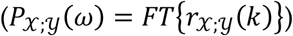, where *ω* ∈ [−π, π] is the normalized angular frequency 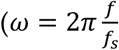 with 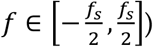. Specifically, the magnitude squared coherence is defined as [28]

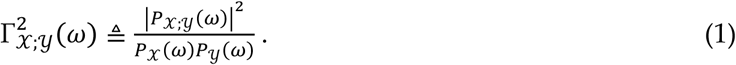

Note that the coherence function is symmetric, 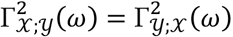, and ranges from 0, indicating that 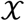 and 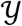 are uncorrelated at the frequency *ω* when 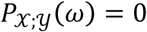, to 1, indicating that 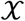 and 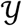 are fully correlated at the frequency *ω* when 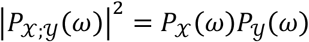.

An alternative, logarithmic measure of linear association in the frequency domain between 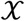 and 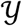 is defined as [29]

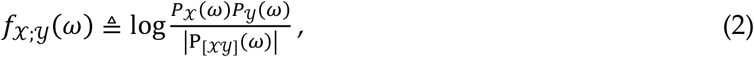

where 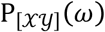 is the 2×2 matrix having the PSDs 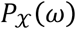 and 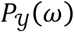 as diagonal elements and the cross-PSDs 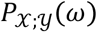 and 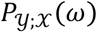 as off-diagonal elements, and |·| stands for matrix determinant. The logarithmic coupling measure (2) can be related to the squared coherence observing that the latter can be formulated from the determinant of the PSD matrix of 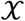 and 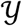 as

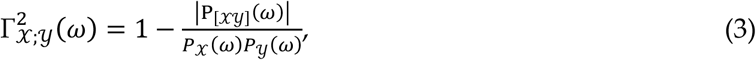

which yields [30]

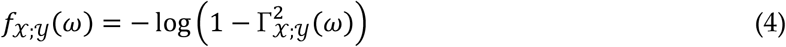

Contrary to the coherence, the logarithmic spectral coupling measure defined in (2) is unbounded as it is null when 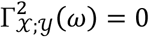 but it tends to infinity when 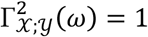. Importantly, this measure has an information-theoretic interpretation, since it has been shown that for Gaussian processes it results as the constituent term of the spectral decomposition of the mutual information rate (MIR) [31,32]. The MIR between 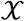 and 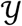 is defined as [31]

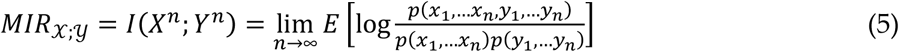

where the vectors *X*^*n*^ = [*X*_1_*X*_2_ … *X*_*n*_] and *Y*^*n*^ = [*Y*_1_*Y*_2_ … *Y*_*n*_] collect the whole dynamics of the two processes observed up to time *n*, and *p*(*x*_1_, … *x*_*n*_) is the joint probability that the variables *X*_1_, …, *X*_*n*_ take the values *x*_1_, …, *x*_*n*_ at the times 1, …, *n* (the same holds for *Y*). The link between frequency and information coupling measures is established by the following relation, which holds when the processes 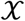 and 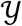 have a joint Gaussian distribution [31,32]:

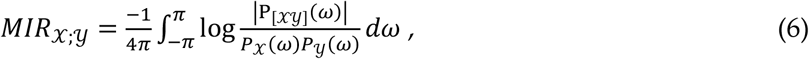

which evidences how the MIR is retrieved integrating the quantity 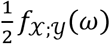 over the whole frequency spectrum. A similar decomposition was proposed by Geweke [33], who introduced the time domain measure of linear dependence

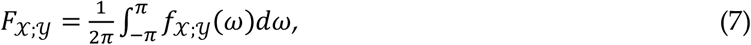

which obtains an information-theoretic interpretation considering (6), i.e., 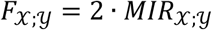. Given this interpretation, and using the natural logarithm in eqs. (2) and (4–6), the quantity 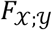 is measured in natural units (nats), and the spectral quantity 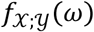 is measured in nats/rad.

### 2.2 Spectral Information Decomposition in Multivariate Processes

Let us consider a multivariate (vector) stochastic process composed by three zero-mean processes 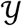, 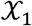, 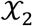 where 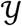 is assumed as “target” and 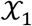 and 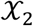 are assumed as “sources”; here the individual processes are taken as scalar, but our formulations extend intuitively to the vector case. To study the interactions among the three processes in the frequency domain, we consider the 3 × 3 spectral density matrix

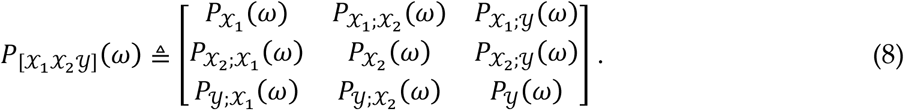

Using this matrix, spectral and information measures of the coupling between each source and the target, i.e. between 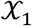 and 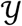 or between 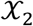 and 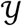, are obtained from the auto- and cross-PSDs as in (2) and (7). Moreover, measures of the coupling between the two sources taken together and the target, i.e. between 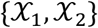 and 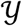, are defined extending (2) and (7) to the multivariate case as

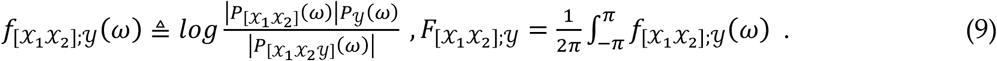

Then, following the principles whereby interaction information is defined for random variables [21,34], we define the following information-theoretic and spectral measures of source interaction:

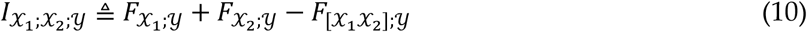

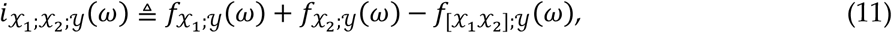

The two measures satisfy the properties of interaction information stated in the time or frequency domains. Specifically, if the two sources 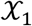 and 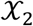 exhibit a stronger additive degree of linear dependence with the target 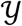 when they are considered separately than when they are considered together 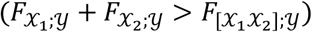, the time-domain source interaction measure is positive 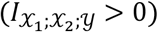, denoting *redundancy*; if, on the contrary, the linear association between 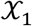 and 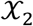 considered jointly and 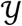 is stronger than the sum of the linear association between 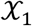 and 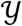 and 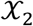 and 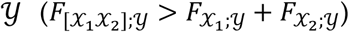, the time-domain source interaction measure is negative 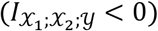, denoting *synergy*. The same properties are satisfied by the frequency domain source interaction measure, and hold for each specific frequency (i.e., redundant and synergistic interaction occur between 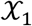 and 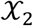 at the frequency *ω* respectively when 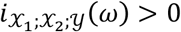 and 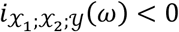. Moreover, thanks to the linearity of the integral operator, the time and frequency domain interaction measures satisfy the property of spectral integration, i.e., 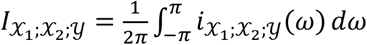.

The decompositions in (10) and (11) provide interaction measures which can take either positive or negative values, denoting respectively redundancy and synergy. An alternative decomposition, which is currently under development in the field of information theory, is the so-called partial information decomposition (PID) [22]. With this decomposition, redundancy and synergy are quantified separately as positive quantities, according to an expansion of the overall interaction between the target and the two source processes that includes also “unique” contributions of each source to the target. Following the philosophy of PID, we define the frequency-specific decomposition of the coupling between 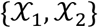 and 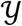 as

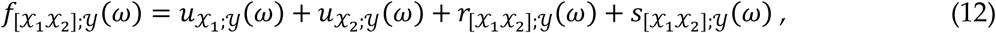

where, at each angular frequency *ω*, 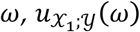 and 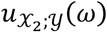 quantify the unique interaction between each source and the target, and 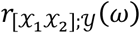 and 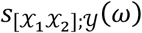 quantify the redundant and synergistic interaction between the two sources and the target. The measures in (12) are defined in a way such that the sum of the unique and redundant interaction between one source and the target yields the corresponding frequency domain coupling measure, i.e.,

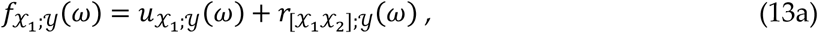

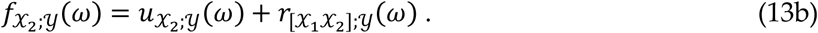

Moreover, a fourth defining equation is needed to compute the PID measures unequivocally; here, exploiting the derivations obtained for Gaussian processes where the same linear representation of process interactions considered in this work is known to completely describe the dynamics [19], we define redundancy as the minimum of the interaction between each individual source and the target, i.e., 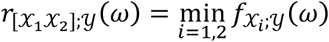[35]. This choice, together with (12) and (13a,b) sets a system of four equations in the four unknowns 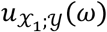, 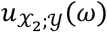, 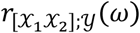, and 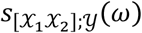, which can thus be computed at each frequency from the three coupling measures 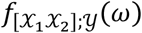, 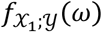 and 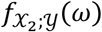.

We note that the PID measures of redundancy and synergy are related to the interaction measure (11). Indeed, substituting (12) and (13) into (11) we obtain 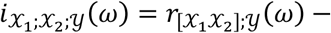 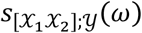, showing that the measure of interaction quantifies the ‘net’ redundancy (i.e., the excess of redundancy over synergy) in the multivariate analysis of the coupling between the two sources and the target. We note also that the frequency-specific PID measures defined above can be integrated to yield equivalent information-rate measures quantifying the unique information shared between 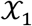 and 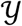 and 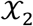 and 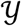, 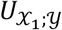 and 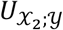, and the redundant and synergistic interaction between 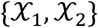 and 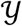, which satisfy (12) in the time domain: 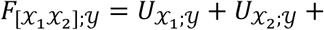 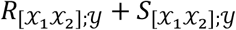. This shows how (12) achieves a spectral decomposition of a time domain PID based on mutual information rates. However, it should be remarked that defining redundancy as the minimum interaction at each frequency prevents from recovering the same property in the time domain, i.e. 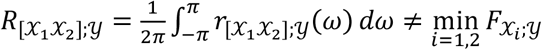.

## 3 Theoretical Example

To illustrate the methodology implemented for the evaluation of frequency domain multivariate interactions among coupled systems, we consider a theoretical example of three Gaussian systems whose associated processes are described by the vector autoregressive (VAR) model with equations

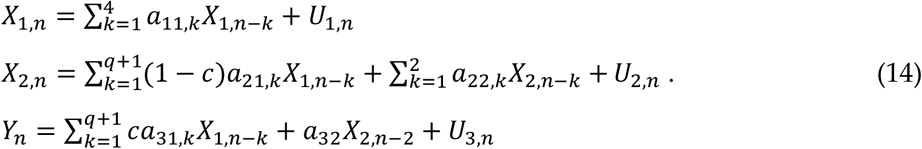

In (14), the simulated source processes 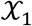 and 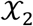 and the target process 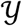 are generated through linear filtering of the innovation processes 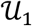, 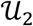, 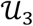, which are set to be white Gaussian noise processes with zero mean and unit variance. The filters are set in a way such that 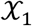 and 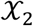 exhibit autonomous stochastic oscillations whose main frequency and bandwidth are modulated by the parameters *a*_11,*k*_ and *a*_22,*k*_, and causal influences are imposed from 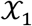 to 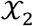, from 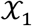 to 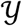 and from 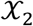 to 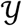 respectively through the coefficients (1 − *c*)*a*_21,*k*_, *ca*_31,*k*_ and *a*_32_ = 1 (see, e.g., [36] for examples of simulated VAR processes). The multiplicative coefficient *c* is a free parameter that is varied to modulate the overall strength of the connection from 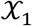 to 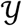 and the inverse overall strength of the connection from 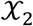 to 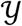. The coefficients *a*_11,1_, …, *a*_11,4_ are set to place two pairs of complex-conjugate poles in the complex plane, with modulus 0.85 and 0.9 and phases 2π · 0.1 rad and 2π · 0.4 rad; similarly, the coefficients *a*_22,1_, *a*_22,2_ are set to place one pair of complex conjugate poles with modulus 0.85 and phase 2π · 0.1 rad. This determines oscillations at the normalized frequency *f*⁄*f*_*s*_ = 0.1 Hz in the PSDs of 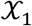 and 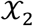 and at the frequency *f*⁄*f*_*s*_ = 0.4 Hz in the PSD of 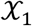 (here we denote frequencies in Hz assuming *f*_*s*_ = 1). The coefficients *a*_21,*k*_, *k* = 1, …, *q* + 1, are obtained as the parameters of a finite impulse response (FIR) filter of order *q* = 16, designed in the high-pass configuration with cutoff frequency of 0.25 Hz; the coefficients *a*_31,*k*_ are set to obtain a low-pass filter of the same order and with the same cutoff frequency. The overall simulation design, depicting autonomous oscillations and the corresponding spectral content, as well as causal connections and the corresponding transfer functions, is depicted in Fig. 1(a).

**Figure 1.**
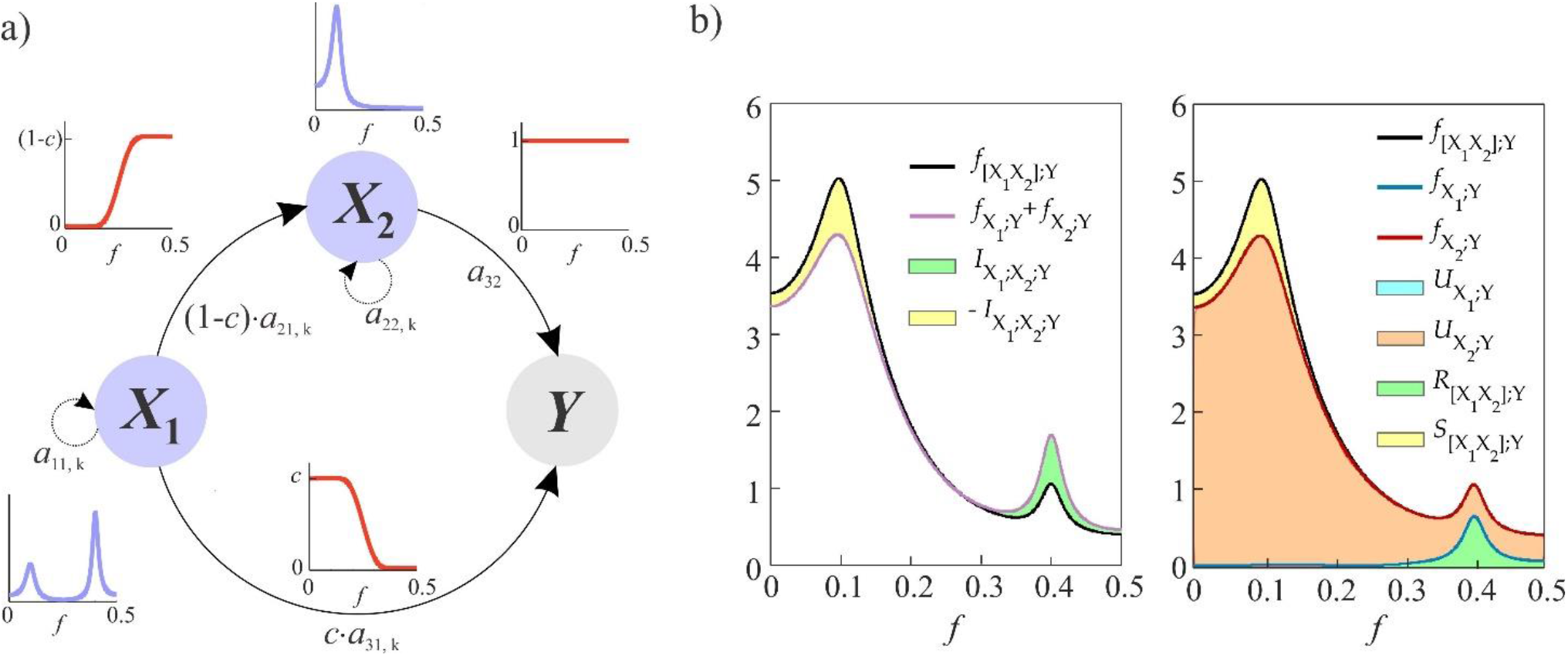
Simulation of coupled linear stochastic processes and exemplary computation of frequency-domain interactions. (a) Schematic representation of the simulated process 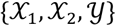 evidencing the autonomous oscillations generated in the sources 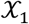 and 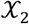 (spectral densities in violet) through the coefficients *a*_11,1_, …, *a*_11,4_ and *a*_22,1_, *a*_22,2_, and the transfer functions (red curves) transmitting such oscillations from 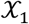 to 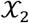 (high-pass, amplitude-modulated by 1 − *c*), from 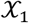 to the target 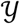 (low-pass, amplitude-modulated by *c*), and from 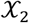 to 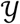 (all-pass). (b) Spectral interaction measures computed for *c* = 0.5, evidencing that the coupling between target and sources peaks at 0.1 Hz (black and red curves) and at 0.4 Hz (black, red, blue curves), that synergy (yellow area) and redundancy (green area) are generated at the two frequencies, and that only the second source provides unique contribution to the coupling (orange area).

The VAR process described above can be studied in the frequency domain by first taking the Z-transform of (14) to yield 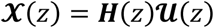, where 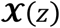 and 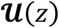 are the Z-transforms of the analyzed processes 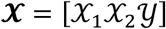 and of the innovations 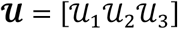, and ***H***(*z*) is the 3 × 3 transfer matrix, then computing the transfer function on the unit circle in the complex plane (***H***(*ω*) = ***H***(*z*)|*z* = *e*^*jω*^), and finally deriving the 3 × 3 spectral density matrix as 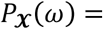 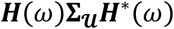 (where 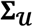 is the covariance of 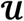 (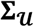 is the identity matrix in our simulation) and * stands for complex conjugate) [27]. This derivation allows to compute the theoretical profiles of the auto and cross-spectral densities contained in 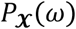 (see also (8)) analytically from the VAR coefficients set in (14). In turn, this leads to compute the exact values of all the time and frequency domain information measures defined in Sect. 2 for the theoretical process, allowing to investigate their behavior given the simulated oscillations and network structure and as a function of the coupling parameter *c*. An example of the analysis for *c* = 0.5 is reported in Fig. 1(b). The left panel shows how interaction information can be elicited at each frequency comparing the joint coupling 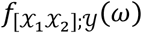 with the sum of the two individual couplings 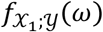 and 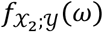: when the joint coupling prevails, negative interaction denoting net synergy is detectable (yellow area around 0.1 Hz); when the individual couplings prevail, positive interaction denoting net redundancy is detectable (green area around 0.4 Hz). The right panel reports the frequency domain PID, showing how redundancy is retrieved as the area below the minimum between the two coupling functions (green, in this case always 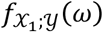, unique contributions are retrieved as the areas between the two coupling functions (in this case only 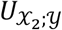 in orange, while 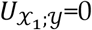), and synergy results as the area depicted in yellow between the joint coupling and the larger of the two individual couplings (in this case always 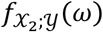).

The results of information decomposition performed in the time and frequency domains for the analyzed VAR process are shown in Fig. 2(a) and Fig. 2(b), respectively. When the coupling parameter *c* is increased from 0 to 1, the simulated interactions change in a way such that the causal coupling from the first source 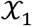 to the target 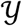 shift progressively towards a smaller importance of indirect interactions (i.e., interactions mediated by the second source 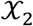) and a larger importance of direct (non-mediated) interactions. This results in a weakening of the interactions from the sources to the target measured by the decrease of the time-domain coupling measures 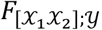, 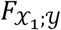 and 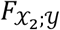 (Fig. 2(a)). More importantly, the reorganization of the interaction structure with the coupling parameter alters the balance between redundancy and synergy: when *c* = 0, the condition of fully indirect effects 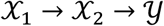 is reflected by positive interaction information 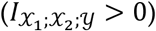 and totally redundant interactions 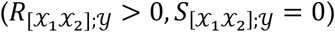; when *c* = 1, the condition of fully independent direct effects 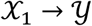 and 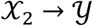 yields negative interaction information with strong prevalence of synergy over redundancy 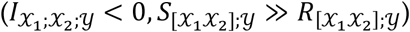. We note also that, since in this simulated network the interaction between 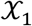 and 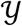 is always weaker than that between 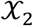 and 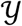, the unique interaction with the target is always ascribed to the second source 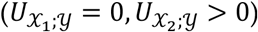.

**Figure 2.**
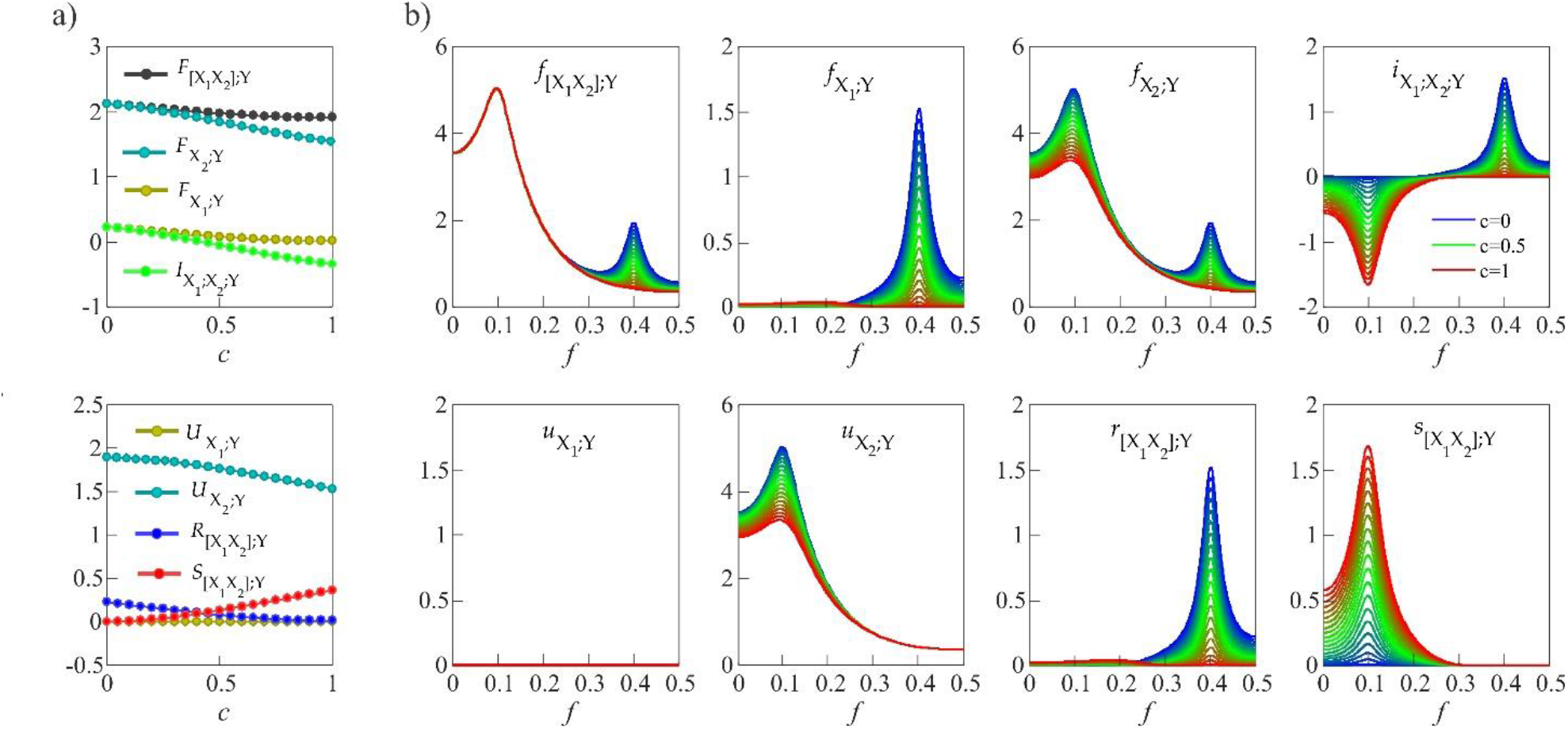
Computation of time- and frequency-domain multivariate information measures for the simulated VAR process depicted in Fig. 1. (a) Time-domain measures of information shared by the two sources and the target jointly 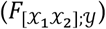, individually 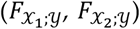 and uniquely 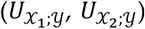, and of interaction information 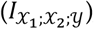, redundant information 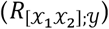 and synergistic information 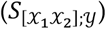 plotted as a function of the coupling parameter *c*. (b) Spectral measures that decompose in the frequency domain the eight time-domain measures (spectral measures are denoted with lowercase letters), plotted as a function of the normalized frequency *f*/*f*_*s*_ for values of the coupling parameter ranging from the case of common mechanism 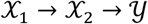 (*c*=0, blue) to the case of separate mechanisms 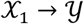 and 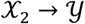 (*c*=1, red), passing by the case of balanced common and separate mechanisms (*c*=0.5, green).

The decomposition of the information measures performed in the frequency domain allows to evidence interaction mechanisms which are specific of the oscillations simulated in the bands centered at 0.1 Hz and 0.4 Hz. The simulation is designed to generate oscillations at 0.1 Hz independently in the two sources 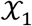 and 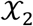, and to couple them separately with the target through the links 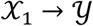 and 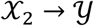; on the contrary, the oscillation at 0.4 Hz generated in 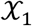 is coupled to the target only indirectly through the chain 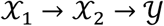. These frequency-specific pathways of interaction are obtained thanks to the low- and high-pass filters that block the transmission of high-frequency oscillations along the direction 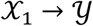, and of low-frequency oscillations along the direction 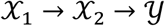, respectively (see Fig. 1(a)), and determine the spectral patterns of information decomposition depicted in Fig. 2(b). The interaction measures peak at the coupling frequencies with a behavior dependent on the coupling parameter: when *c* is low, the direct transmission of the slow oscillation from the second source to the target through the link 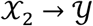 is revealed by the peak at *ω* = 2π0.1 rad in 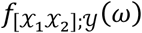 and 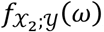, while indirect transmission of the fast oscillation the chain 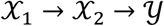 is revealed by the peaks at *ω* = 2π0.4 rad in all the three interaction measures; when *c* increases towards 1, the 0.1 Hz oscillation is transmitted independently from each source to the target still determining low-frequency peaks in the interaction measures, while the 0.4 Hz oscillation is blocked (by the coefficient 1 − *c* along 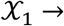 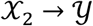 and by the low-pass filter along 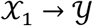) so that the high-frequency peaks in 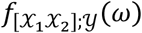 and 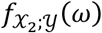 vanish progressively at increasing *c*.

As regards the interaction information and PID measures, their frequency decomposition reveals the nature and the strength of the interaction mechanisms simulated within specific frequency bands. In particular, when only the indirect coupling 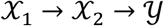 was simulated (*c* = 0, blue lines in Fig. 2(b)), the interaction information shows a positive peak at 0.4 Hz which corresponds to full redundancy 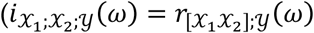, 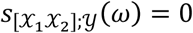 for *ω* = 2π0.4), while all three measures are null at 0.1 Hz; when only the two indirect couplings 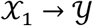 and 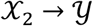 were simulated (*c* = 1, red lines in Fig. 2(b)), the interaction information shows a negative peak at 0.1 Hz which corresponds to full synergy 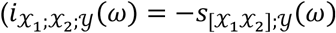, 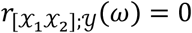 for *ω* = 2π0.1), while all three measures are null at 0.4 Hz; in the intermediate case when both direct and indirect coupling mechanisms were active (*c* = 0.5, green lines in Fig. 2(b), corresponding to the case depicted in Fig. 1(b)), the interaction information peaks both at 0.1 Hz and at 0.4 Hz showing respectively full synergy and full redundancy 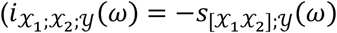, 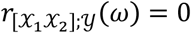 at *ω* = 2π0.1;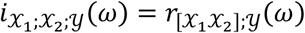 at *ω* = 2π0.4). This latter situation corresponds to the interesting case of coexistence of fully synergetic and fully redundant mechanisms operating in different frequency bands for the same three interacting signals. We note that, while the coexistence of synergy and redundancy with *c* = 0.5 is evident looking at the spectral profile of the interaction information 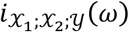, it is not observable when only the time domain measure 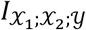 is computed.

The simulation reported in this section illustrates the meaning of multivariate information measures computed in both time and frequency domains between three interacting signals. We have shown how indirect coupling effects (coupling between one source and the target mediated by the other source) are reflected by positive interaction information and prevalence of redundancy, while direct coupling effects occurring independently between each source and the target are reflected by negative interaction information and prevalence of synergy. Moreover, the spectral decomposition of the interaction measures allows to ascribe the observed coupling mechanisms to oscillatory behavior of the analyzed signals: we have reported instances of fully redundant and fully synergistic interactions, occurring either in an exclusive way at specific frequencies or even simultaneously in different frequency bands. Importantly, the use of frequency domain measures can elicit interaction mechanisms which are masked in the time domain, such as the simultaneous presence of synergistic and redundant coupling revealed when interaction information is integrated in distinct frequency bands but not when it is integrated over the whole frequency axis.

## 4 Application to Cardiovascular and Cardiorespiratory Interactions

This section reports the application of the framework for the analysis of multivariate interactions in the frequency domain to an historical database of short-term cardiovascular variability series previously studied in the context of information decomposition [13,16,25,26]. The present analysis is performed on the beat-to-beat time series of heart period, systolic and diastolic arterial pressure to study cardiovascular interactions, and on the series of heart period, systolic pressure and respiration to study cardiorespiratory interactions. Spectral analysis is focused on the two frequency bands of physiological interest, i.e. the low frequency (LF, 0.04-0.15 Hz) and high frequency (HF, 0.15-0.4 Hz) bands, to illustrate how distinct physiological mechanisms operating within these bands can be described through the computation of frequency-specific measures of coupling, interaction information and redundancy/synergy.

### 4.1 Experimental Protocol and Data Analysis

We consider sixty-one healthy volunteers (37 females, 24 males, 17.5 ± 2.4 years old) free of medications or substances influencing autonomic nervous system or cardiovascular system activity, who were enrolled in a study aimed at monitoring cardiovascular variability during physiological stress [6,13]. All procedures were approved by the Ethical Committee of the Jessenius Faculty of Medicine, Comenius University, Martin, Slovakia, and all participants gave written informed consent. Here we study data acquired during a resting state in the supine body position (condition R) and during passive head-up tilt in the 45 degrees body position (condition T). Despite the small magnitude of the orthostatic challenge, during 45 degrees T evident signs of vagal withdrawal are detectable as suggested by the significant decrease of respiratory sinus arrhythmia and a decrease of baroreflex sensitivity [12] as well as signs of sympathetic activation as denoted by a remarkable increase of QT variability [37].

The acquired signals were the continuous finger arterial blood pressure recorded non-invasively by the photoplethysmographic volume-clamp method (Finometer Pro, FMS, Netherlands), the electrocardiogram (horizontal bipolar thoracic lead; CardioFax ECG-9620, NihonKohden, Japan) and the respiratory volume recorded by respiratory inductive plethysmography using thoracic and abdominal elastic bands (RespiTrace, NIMS, USA). From these signals, four time series were measured on a beat-to-beat basis to quantify the spontaneous variability of the basic cardiovascular variables together with respiration: the heart period series (*H*) was obtained as the sequence of the temporal distances between consecutive R waves of the ECG (R-R intervals), the systolic and diastolic blood pressure series (*S*, *D*) were obtained as the sequences of the maximum and minimum values of the arterial blood pressure signal measured within each detected R-R interval, and the series of respiration values (*R*) was obtained as the sequence of respiratory volume values sampled at the onset of each detected R-R interval. For each subject and experimental condition, stationary segments of *N*=300 points were selected synchronously for the four series (an example is reported in Fig. 3(a)). Before the analysis, the series were linearly de-trended removing the best straight-fit line and were then reduced to zero mean. The resulting time series were considered as realizations of the stochastic processes describing the dynamics of the heart period, systolic and diastolic pressure, and respiration (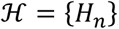, 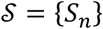, 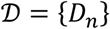, 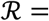 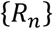, *n* = 1, …, *N*).

**Figure 3.**
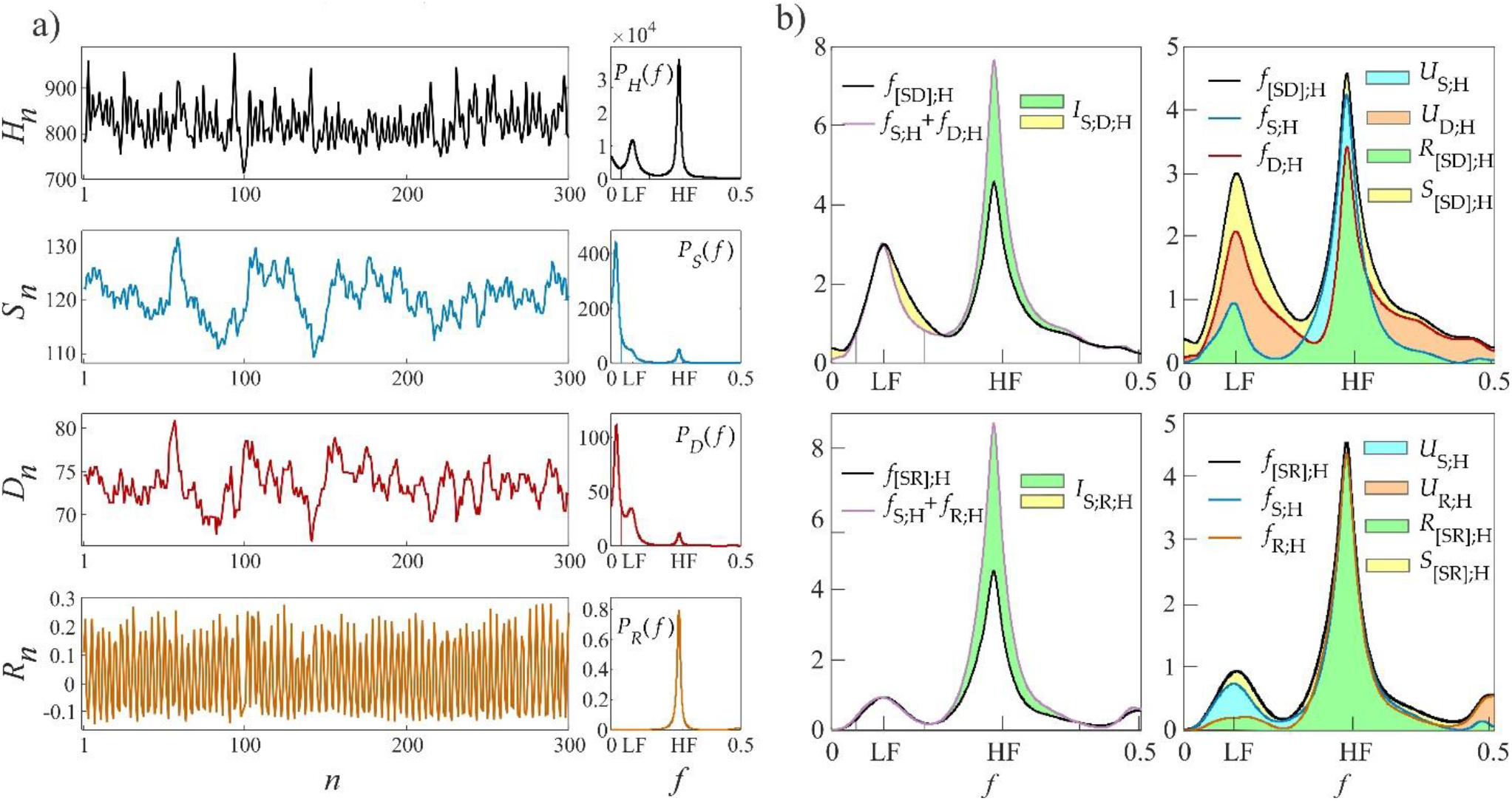
Example of frequency domain information-theoretic analysis of cardiovascular and cardiorespiratory multivariate interactions. (a) Time series of heart period (*H*), systolic pressure (*S*), diastolic pressure (*D*) and respiration volume (*R*) measured in the resting supine position for a representative subject; the power spectral density estimated for each time series are depicted on the right, with vertical lines delimiting the LF band (0.04-0.15 Hz) and the HF band (0.15-0.4 Hz). (b) Spectral profiles of the information measures computed taking systolic and diastolic pressure as sources and heart period as target process (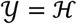, 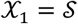, 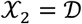, upper panels), and taking systolic pressure and respiration as sources and heart period as target process (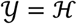, 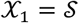, 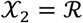, lower panels). In the two cases, the left panels depict the computation of the interaction information 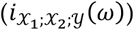 as the difference between the information shared by the target and the two sources individually 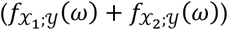 and jointly (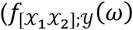, black curve), with integrated contributions resulting as redundant (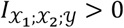, green areas) or synergistic (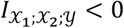, yellow areas); the right panels depict the frequency domain PID, showing how redundancy (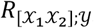, green areas) emerges as the information integrated by the minimum coupling function between each source and the target (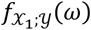 and 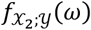, blue and red/orange curves), unique information (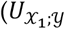 or 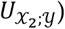) is integrated between the two coupling functions (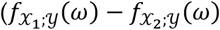, light blue areas, or 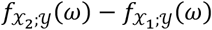, light orange areas), and synergy (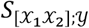, yellow areas) results as the information integrated between the highest coupling function and the joint coupling function. Band-specific values are obtained for each function limiting the integration within the LF and HF bands.

The analysis was performed considering two separate settings: to study cardiovascular interactions, the heart period was considered as the target process and the systolic and diastolic pressure were considered as sources, i.e., 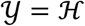, 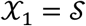, 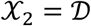; to study cardiorespiratory interactions, the target and first source were left unchanged and the second source was taken to be the respiratory volume, i.e., 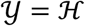, 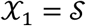, 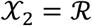. For each of the two settings, and for each subject and experimental condition, parametric estimates of spectral information measures were obtained as follows. First, a VAR model was identified on the three time series, using the ordinary least squares method to estimate the model coefficients and setting the model order according to the Akaike Information Criterion [27]. Then, the estimated model parameters (VAR coefficients and covariance matrix of the prediction errors) were used to yield estimates of the PSD matrix for the three processes according to (8) and as described in Sect. 3. Finally, estimates of the frequency-domain functions measuring the information shared between the target and the two sources taken separately (eq. (2)) or jointly (eq. (9)), the interaction information between target and sources (eq. (10)), as well as the unique, redundant and synergistic terms of the PID of target-source interactions (eq. (12)), were obtained from the estimated PSDs.

All spectral measures were computed for frequencies ranging from 0 to the Nyquist frequency *f*_*s*_/2, where the sampling frequency *f*_*s*_ was assumed as the inverse of the mean RR interval. For each spectral information measure, values indicative of the overall information shared among the processes, and of the information shared in the LF and HF bands, were obtained by integration of the measure over the appropriate frequency ranges and multiplication by a normalization factor properly set to satisfy (7); for instance, values of the interaction information quantified accounting for the whole frequency range, for the LF range, and for the HF range, are obtained respectively as 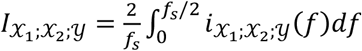, 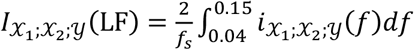, and 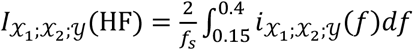. An example of computation of the spectral information measures and of their evaluation within the LF and HF bands is reported in Fig. 3. The analysis for this subject shows well defined power content (Fig. 3(a)) in the LF and HF bands for 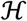, 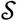 and 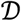, corresponding to LF and HF peaks for the coupling measures 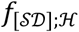, 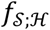 and 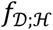 (Fig. 3(b), upper panels), and in the HF band for 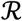, corresponding to a dominant HF peak for the coupling measures 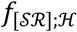, 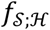 and 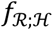 (Fig. 3(b), lower panels); the cardiovascular interaction information 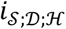 is negative in the LF band and positive in the HF band, reflecting nontrivial amounts of synergy 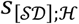 in LF and of redundancy 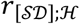 in HF (respectively, yellow and green areas in Fig. 3(b), upper panels), while the cardiorespiratory interaction information 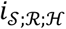 is high and positive in the HF band, reflecting marked f redundancy 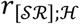 in HF (green areas in Fig. 3(b), lower panels).

### 4.2 Results and Discussion

Results of the spectral analysis of information measures are reported in Figs. (4) and (5) for the study of cardiovascular interactions (setting 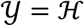, 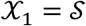, 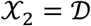) and in Figs. (6) and (7) for the study of cardiorespiratory interactions (setting 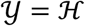, 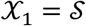, 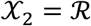). Results are presented showing the distributions (expressed as boxplots and individual values) of the spectral interaction measures (Figs. 4,6) and of the PID measures (Figs. 5,7) computed at rest (R) and during head-up tilt (T) and integrated over the whole frequency range (TOT) or within the LF or HF bands. For each measure and frequency band, the distributions measured during R and T are compared with a two-sided Wilcoxon signed rank test for paired data in order to assess the statistical significance of the changes induced in the measure by the orthostatic stress; a *p*-value <0.05 was considered as statistically significant for each comparison.

**Figure 4.**
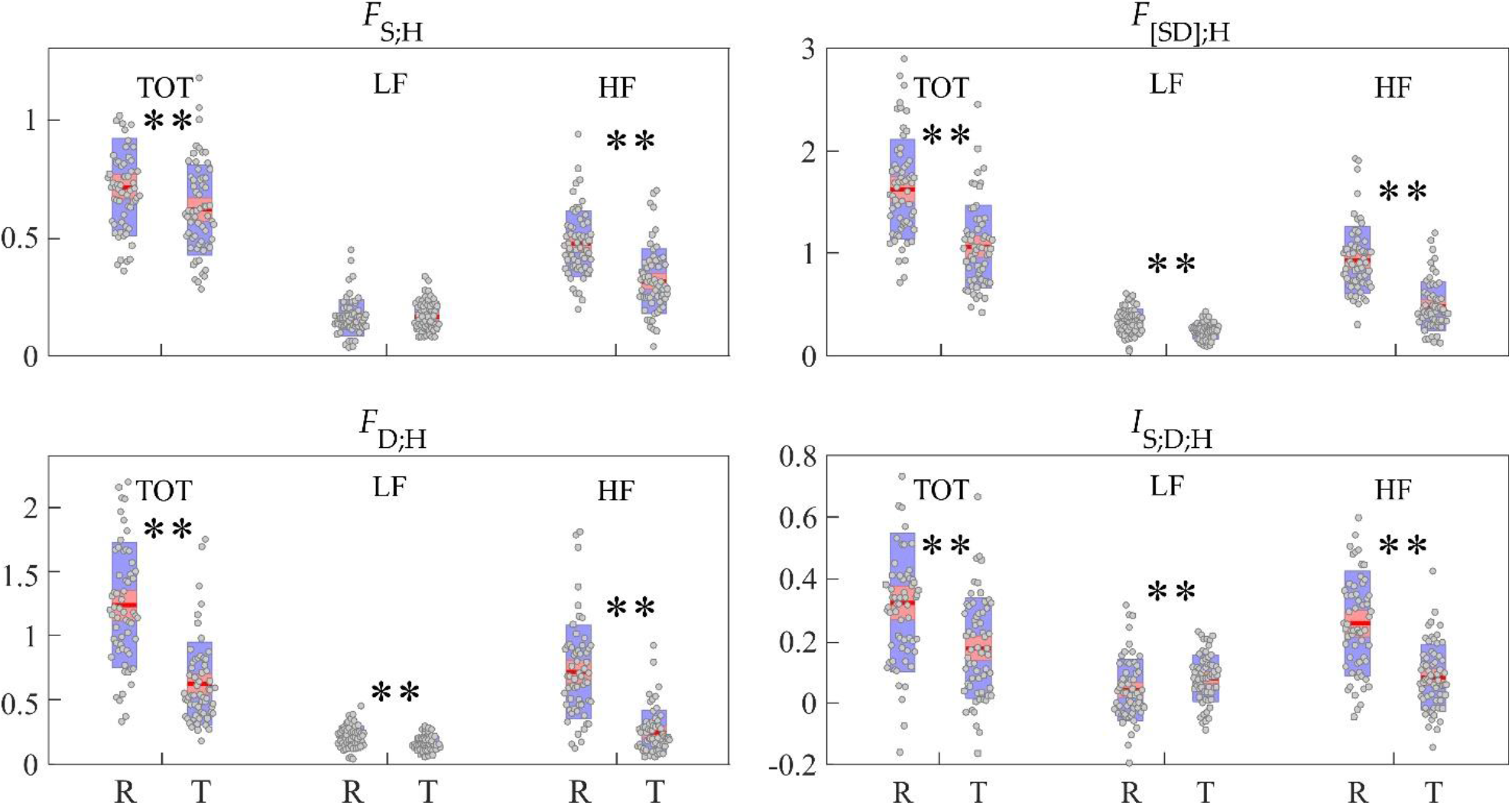
Spectral information decomposition of multivariate cardiovascular interactions (target: heart period 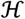; sources: systolic pressure 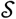, diastolic pressure 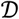). Plots depict the distribution across subjects of the information shared by the target and the two sources jointly 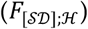 and individually (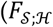, 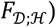), and of the interaction information between the two sources and the target 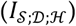 obtained integrating the corresponding spectral measures over the whole frequency range (TOT), in the range 0.04-0.15 Hz (LF) and in the range 0.15-0.4 Hz (HF) and computed in the resting state (R) and during head-up tilt (T). Wilcoxon test comparing R and T: *, p<0.05; **, p<0.005.

**Figure 5.**
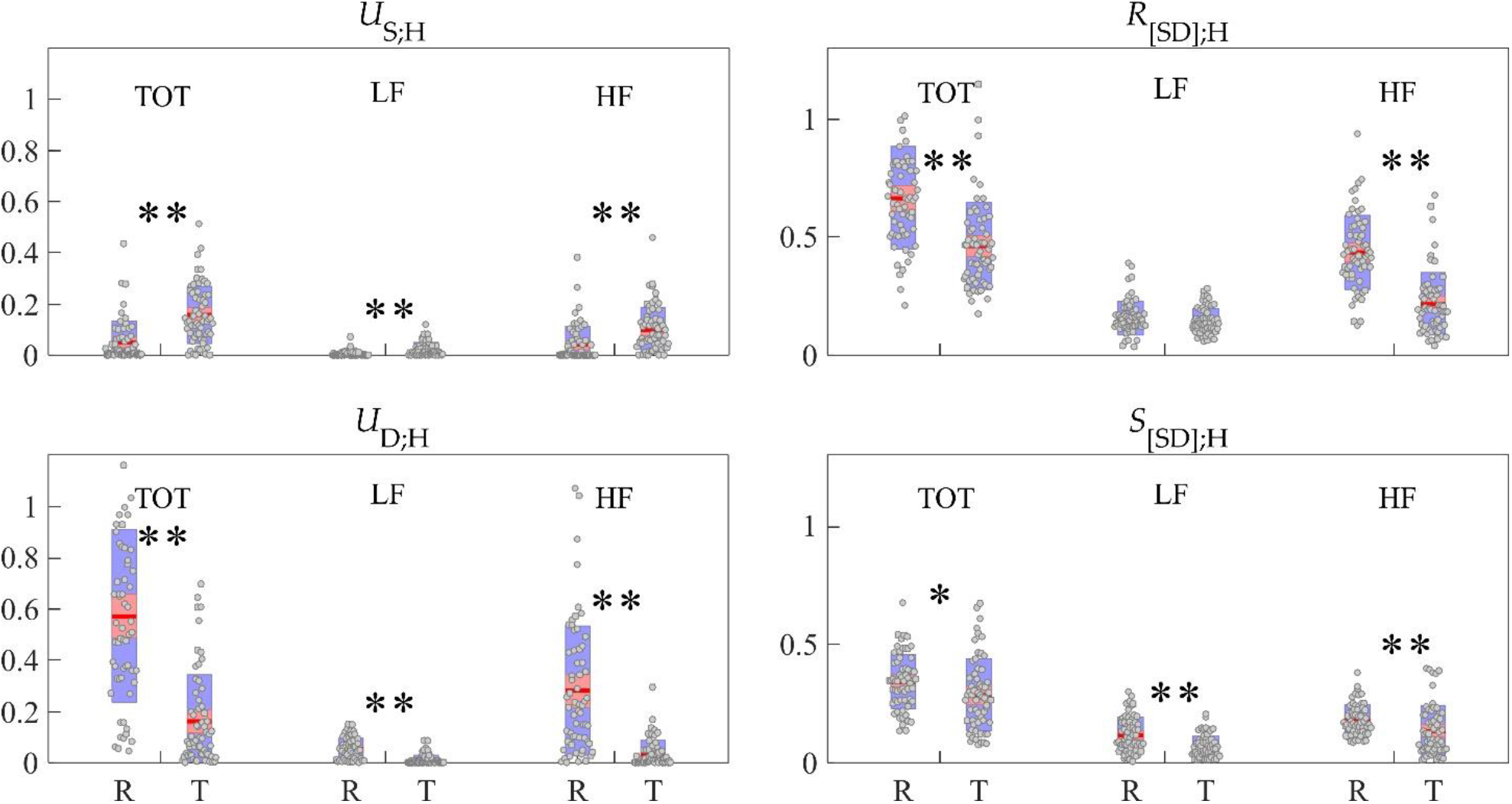
Spectral partial information decomposition of multivariate cardiovascular interactions (target: heart period 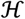; sources: systolic pressure 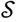, diastolic pressure 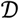). Plots depict the distribution across subjects of the unique information shared by the target and each source (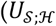, 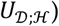), and of the redundant 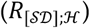 and synergistic 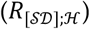 information between the two sources and the target obtained integrating the corresponding spectral measures over the whole frequency range (TOT), in the range 0.04-0.15 Hz (LF) and in the range 0.15-0.4 Hz (HF) and computed in the resting state (R) and during head-up tilt (T). Wilcoxon test comparing R and T: *, p<0.05; **, p<0.005.

**Figure 6.**
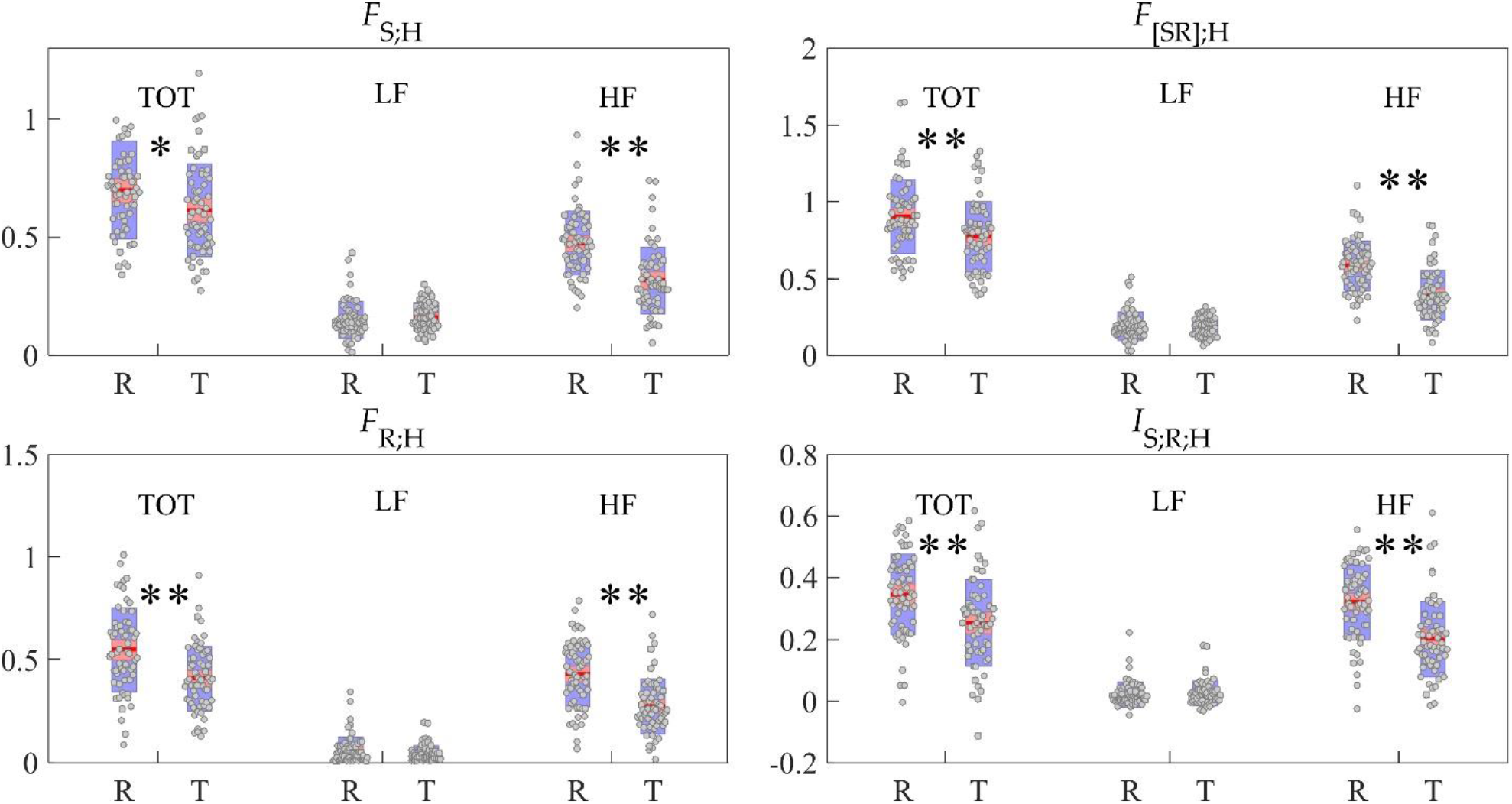
Spectral information decomposition of multivariate cardiorespiratory interactions (target: heart period 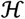; sources: systolic pressure 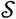, respiration 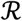). Plots depict the distribution across subjects of the information shared by the target and the two sources jointly 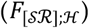 and individually (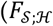, 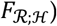), and of the interaction information between the two sources and the target 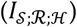 obtained integrating the corresponding spectral measures over the whole frequency range (TOT), in the range 0.04-0.15 Hz (LF) and in the range 0.15-0.4 Hz (HF) and computed in the resting state (R) and during head-up tilt (T). Wilcoxon test comparing R and T: *, p<0.05; **, p<0.005.

**Figure 7.**
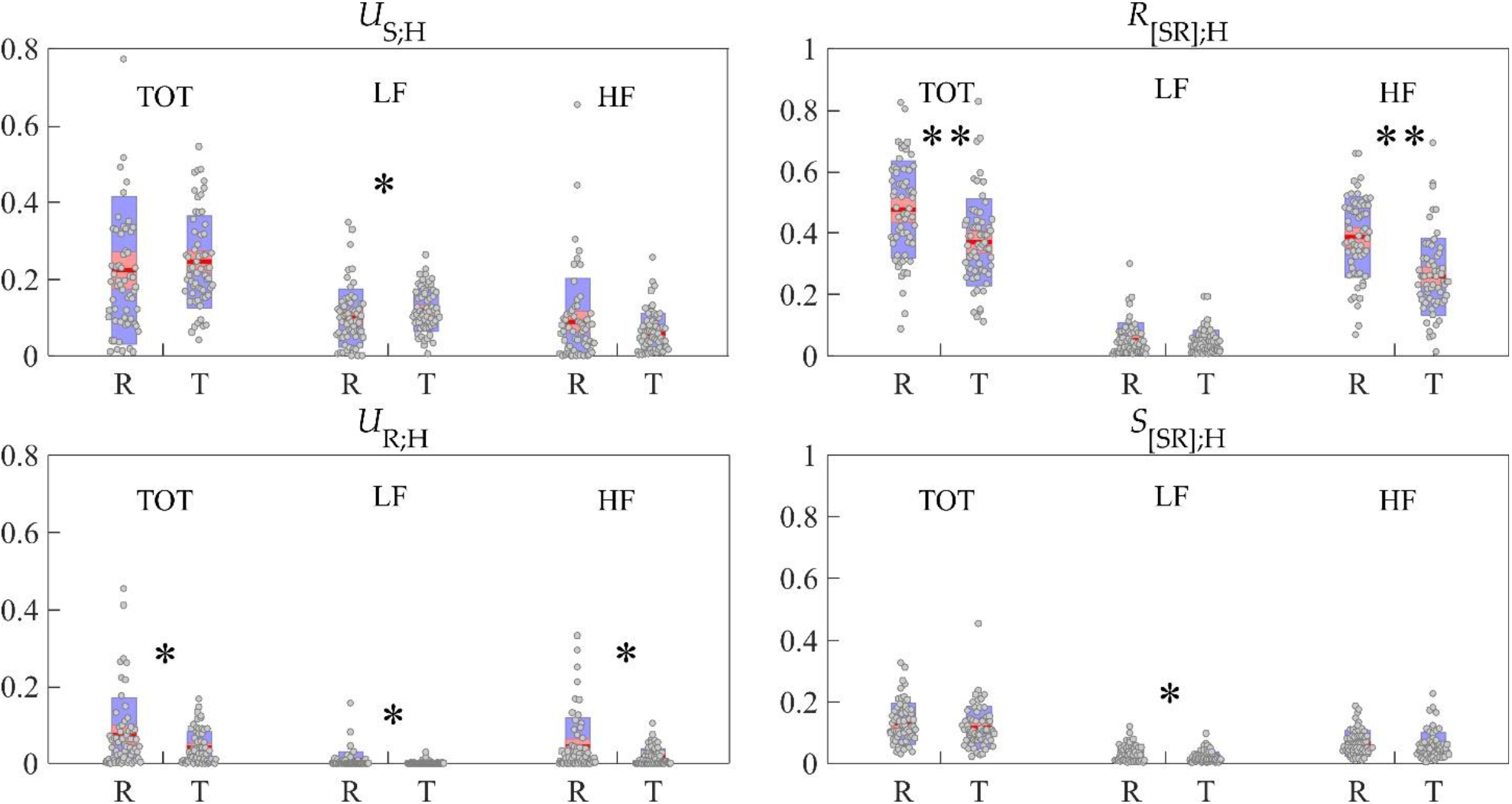
Spectral partial information decomposition of multivariate cardiorespiratory interactions (target: heart period 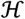; sources: systolic pressure 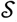, respiration 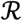). Plots depict the distribution across subjects of the unique information shared by the target and each source (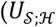, 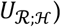), and of the redundant 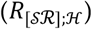 and synergistic 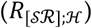 information between the two sources and the target obtained integrating the corresponding spectral measures over the whole frequency range (TOT), in the range 0.04-0.15 Hz (LF) and in the range 0.15-0.4 Hz (HF) and computed in the resting state (R) and during head-up tilt (T). Wilcoxon test comparing R and T: *, p<0.05; **, p<0.005.

The analysis of heart period, systolic pressure and diastolic pressure dynamics indicates the presence of strong cardiovascular interactions which occur mainly in the HF band of the frequency spectrum and tend to decrease with the transition from the supine to the upright body position. Indeed, Fig.4 shows high values (generally between 0.5 and 2 nats) of the information shared between the cardiac and the two vascular processes taken together 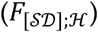 or individually 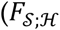, 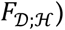, with higher values for the measures integrated in the HF band, and statistically significant decreases from R to T of 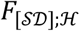 and 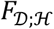 (both over the whole spectrum (TOT) and within LF and HF bands) and of 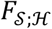 (TOT and HF values). On the other hand, the application of PID (Fig. 5) reveals that, in comparison with rest, the unique information shared between the heart period and blood pressure dynamics during tilt is lower considering the diastolic pressure (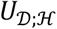 decreases significantly from R to T in all bands) but is actually higher considering the systolic pressure (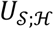 increases significantly from R to T in all bands). Redundant interactions between systolic and diastolic pressure dynamics are prevalent over synergistic interactions, as documented by the mostly positive values of 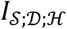 in all bands (Fig. 4). Interestingly, moving from R to T the net redundancy increases significantly in the LF band, and decreases significantly in the HF band. Decomposing the net redundancy in separate redundant and synergistic effects shows that 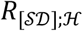 is stable in the LF band and decreases in the HF band, while 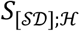 decreases significantly in both bands (Fig. 5). These results about interaction information decomposition suggest that the postural change is associated, in the LF band, with an increased importance of common mechanisms of interaction between systolic/diastolic pressure and heart period (interactions 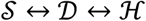), which is due to a weakening of separate mechanisms (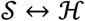 and 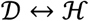); in the HF band, a decreased importance of both common and separate mechanisms of vascular and cardiac interaction is observed.

The high values observed for the measures of cardiovascular coupling can be explained physiologically with reference to several regulatory mechanisms, including the arterial baroreflex that reflects feedback interactions from systolic pressure to heart period (interaction between 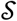 and 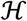), the cardiac runoff that relates the duration of the heart period with ventricular filling and the pressure at the end of the diastole (interaction between 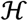 and 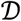), and the Frank-Starling law that is responsible for the relation between the increased end-diastolic volume and strength of the systolic contraction (relation between *H* and 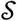) [5,6,38]. Our findings documenting higher strength of the interactions between 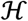 and 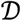 compared with those between 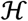 and 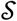 agree with those reported by Javorka et al. [6], who also detected strong directional interactions along the chain 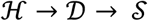 which are compatible with the dominance of redundancy (positive 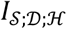 and high 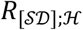) observed in the present study. The findings in [6] include also a decreased strength of such directional interactions during head-up tilt, possibly related to the lower magnitude of heart period oscillations which may have an effect on the cardiac runoff [39], which also agrees with the decreased interaction information observed in this study with the orthostatic challenge. A reduction of the net redundancy of the cardiovascular control during postural stress was observed also by Porta et al. [24], who also showed that net redundancy is under the ANS control as it decreases in proportion to the vagal withdrawal during graded head-up tilt. Here, we document that such a decrease is localized more in the HF band, thus confirming the involvement of parasympathetic withdrawal leading to decreased magnitude of respiration-related heart rate oscillations, in the weakening of the interactions between systolic/diastolic pressure and heart period observed during tilt. The opposite behavior of interaction information observed in the LF band suggests that the cardiovascular response to postural stress is more complex, with a probable involvement of the sympathetic nervous system in the control of the coupling of low-frequency oscillations of heart period systolic/diastolic pressure. Indeed, our results confirm also the relatively low coupling between 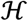 and 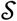 at rest and its increase with head-up tilt observed in previous works [6,9,13,40], documenting the baroreflex response typically associated with sympathetic activation and rise of LF components in the spectra of systolic pressure and heart period.

The analysis performed including the variability of the respiratory volume, depicted in Figs. 6 and 7, indicates that cardiorespiratory interactions are almost exclusively relevant to the HF band of the spectrum. This observation is supported by the small values of the information measures including the respiration process 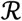 and computed in the LF band, as well as by the same response to tilt observed in all spectral information measures after integration in the whole frequency range or integration in the HF band only. Such a response indicates a general weakening of cardiorespiratory interactions during postural stress, as documented by the statistically significant decrease observed moving from R to T in the information shared both globally and in the HF band between heart period and respiration variability, either individually or jointly with systolic pressure (measures 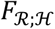 and 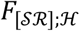, Fig. 6) and even uniquely after PID (measure 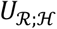, Fig. 7). The transition from rest to tilt induced a significant decrease also in the interaction information 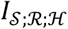, that takes positive values in the large majority of subjects in both conditions (Fig. 6). This finding indicates a prevalence of redundancy over synergy that is confirmed also by PID looking at the small values of synergistic information 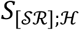, and at the high values of the redundant information 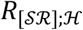, which decreases significantly moving from R to T (Fig. 7). The prevalence of redundancy indicates the importance of common pathways of interaction occurring between 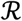 and 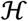 mediated by 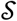. Finally, we note that the computation of measures not involving respiration (i.e. 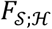 in Fig. 6 and 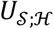 in Fig. 7) leads to similar results to those observed for the same measures computed in the presence of 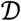 in place of 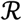 (respectively, Fig. 3 and Fig. 4), showing a tendency of the information shared between heart period and systolic pressure to increase when computed in the LF band, and to decrease when computed in the HF band, with the transition from R to T.

The little relevance within the LF band of the measures of multivariate interaction involving the respiratory volume is due to the known fact that the variability of respiration and its effects on the variability of heart period and arterial pressure are almost exclusively confined in the HF band of the spectrum [2]. The significant decrease with tilt of the information shared between respiration and heart period (measures 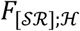, 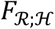, 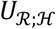) confirms a number of previous investigations based on entropy analysis [10,11,16,25] that documented a reduction of cardiorespiratory interactions during orthostatic stress. Physiologically, these findings reflect the vagal withdrawal and the dampening of respiratory sinus arrhythmia occurring with tilt [39]. Finally, the analysis of the measure of interaction information 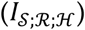 and of its expansion into purely redundant and purely synergistic terms (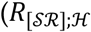, 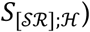) helps in investigating the two physiological mechanisms that may explain the effects of respiration on the heart period known and respiratory sinus arrhythmia (RSA), i.e. the direct pathway whereby neural commands originating in the respiratory centers of the brainstem act on the sinus node independently of blood pressure [8], and the indirect pathway whereby respiratory effects on the arterial pressure are transmitted to the heart period via the baroreflex [7]. In a recent work [26] we have shown how the PID performed defining redundancy as the minimum information contributed by each source to the target [35] ascribes redundancy to the indirect mechanism 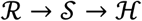, the unique cardiorespiratory transfer to the direct mechanism 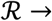 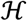, and synergy to the simultaneous activation of both pathways. According to this interpretation, the dominance of redundancy 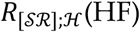(HF) suggests that in our data respiratory sinus arrhythmia is mostly due to the baroreflex-mediated mechanism, and the decrease of both 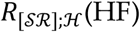(HF) and 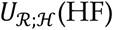(HF) with tilt suggests that the mechanism is dampened with tilt in parallel with the parasympathetic withdrawal. While the prevalence of baroreflex over central mechanisms of RSA was observed also in [26], a different interpretation was given to the response to tilt; we ascribe the difference to the fact that the analysis performed here can be focused on the HF band, and thus allows to exclude more safely the strong variations induced by tilt in the LF band which are otherwise incorporated in non frequency-specific analyses such as that in [26].

## 5 Conclusions

The aim of this study was to introduce a spectral representation of information decomposition able to quantify, within predefined frequency bands associated to specific oscillatory rhythms, the unique, redundant and synergistic information shared by two source processes and one target process describing the activity of a multivariate dynamical system. The proposed framework was designed building upon long-known but not extensively developed relations between information measures and spectral measures valid for Gaussian systems [31,32]. Of note, although the proposed measures were developed on the basis of the linear parametric (VAR) representation of multivariate processes, they can be easily implemented also using non-parametric spectral estimators [28], as they are entirely derived from PSD functions.

The main distinctive feature of the proposed framework is the combination of spectral decomposition and information decomposition, which allows to elicit aspects of the interaction among processes which cannot be observed by information measures alone or by spectral measures alone. The spectral decomposition of information measures allows to retrieve amounts of information shared by the observed processes within specific frequency bands, and thus related to specific oscillatory components. This can be seen, in the theoretical example, when redundant and synergistic amounts of information shared at different frequencies are evident from the positive and negative values of the interaction information measure 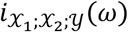 evaluated at those frequencies, but disappear in the integrated time-domain measure 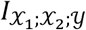. On the other hand, the information decomposition of spectral measures allows to evidence aspects of the coupling between processes that are masked in the analysis using classical frequency domain measures. This occurs in our simulation observing that the coupling between one source and the target is entirely redundant with the other source, so that it appears in the “global” measure 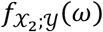 but is completely removed in the measure of unique information 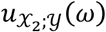 obtained through PID.

The potentiality of combining information and spectral decompositions was verified experimentally in the proposed application to cardiovascular and cardiorespiratory interactions. Indeed, although significant modifications of physiological dynamics induced by the transition from rest to tilt are detectable using simpler univariate variability markers (see, e.g., [12,37,41,42]), the proposed multivariate indexes can describe peculiar features that might be correlated with specific properties of the physiological dynamical systems that go beyond their traditional assessment based on power of oscillations, transfer function gain and latency but deal with higher level functions related to the overall organization of the cardiovascular control system like synergy and redundancy. Specific results observed in this work are the opposite response to tilt observed integrating the cardiovascular interaction information 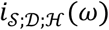 within the LF and HF frequency bands, with the increase in the LF band and the decrease in HF band associated respectively to sympathetic activation and parasympathetic withdrawal, and the different trends observed with tilt for the global and unique measures of coupling between HP ad SAP, with tendency to decrease for 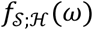 and to increase for 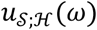 that highlight the dampening of redundant interactions in the response to postural stress.

Future studies are envisaged to extend the proposed framework to dynamic analysis, making it possible to achieve a spectral decomposition of directed information measures like the joint, redundant and synergistic transfer entropy, up to now available only in their time domain [16] and multiscale [18] formulations. From a practical viewpoint, the generalization of the framework to vector processes would allow to describe multi-system interactions in a more exhaustive way, with obvious applications in the fields of computational neuroscience and network physiology [15].

## Additional Information

### Ethics

The experimental part of this study was approved by the Ethical Committee of the Jessenius Faculty of Medicine, Comenius University, Martin, Slovakia, and all participants gave written informed consent.

### Data Accessibility

The data used and the Matlab Software relevant to this work are freely downloadable from www.lucafaes.net/fdPID.html

